# The eyes reflect an internal cognitive state hidden in the population activity of cortical neurons

**DOI:** 10.1101/2020.06.29.178251

**Authors:** Richard Johnston, Adam C. Snyder, Sanjeev B. Khanna, Deepa Issar, Matthew A. Smith

**Author notes:** Corresponding author: Matthew A. Smith, PhD, Carnegie Mellon University, Department of Biomedical Engineering and Neuroscience Institute, Mellon Institute, Room 115, Pittsburgh, PA 15213.

## Abstract

Decades of research have shown that global brain states such as arousal can be indexed by measuring the properties of the eyes. Neural signals from individual neurons, populations of neurons, and field potentials measured throughout much of the brain have been associated with the size of the pupil, small fixational eye movements, and vigor in saccadic eye movements. However, precisely because the eyes have been associated with modulation of neural activity across the brain, and many different kinds of measurements of the eyes have been made across studies, it has been difficult to clearly isolate how internal states affect the behavior of the eyes, and vice versa. Recent work in our laboratory identified a latent dimension of neural activity in macaque visual cortex on the timescale of minutes to tens of minutes. This ‘slow drift’ was associated with perceptual performance on an orientation-change detection task, as well as neural activity in visual and prefrontal cortex (PFC), suggesting it might reflect a shift in a global brain state. This motivated us to ask if the neural signature of this internal state is correlated with the action of the eyes in different behavioral tasks. We recorded from visual cortex (V4) while monkeys performed a change detection task, and the prefrontal cortex, while they performed a memory-guided saccade task. On both tasks, slow drift was associated with a pattern that is indicative of changes in arousal level over time. When pupil size was large, and the subjects were in a heighted state of arousal, microsaccade rate and reaction time decreased while saccade velocity increased. These results show that the action of the eyes is associated with a dominant mode of neural activity that is pervasive and task-independent, and can be accessed in the population activity of neurons across the cortex.

## Introduction

In the fields of psychology and neuroscience, the eyes are often viewed as a window to the brain. Much has been learned about cognitive processes, and their development, from studying the action of the eyes (Aslin, 2012; Eckstein, Guerra-Carrillo, Miller Singley, & Bunge, 2017; Hannula et al., 2010; Hessels & Hooge, 2019; König et al., 2016; Ryan & Shen, 2020). In addition, a large body of research has shown that properties related to the eyes can be used to index global brain states such as arousal, motivation and cognitive effort (Di Stasi, Catena, Cañas, Macknik, & Martinez-Conde, 2013; Joshi & Gold, 2019; Mathôt, 2018; C. A. Wang & Munoz, 2015). The action of the eyes can be considered broadly in two contexts – the action of the pupil when the eyes are relatively stable, and the action of the eyes when they move, be it voluntarily, in response to novel objects in the visual field, or involuntarily during periods of steady fixation. In each context, technological advancements in infrared eye-tracking have allowed rich insight about a subject’s global brain state to be surmised in a rapid, accurate and non-invasive manner (Kimmel, Mammo, & Newsome, 2012).

When the eyes are relatively stable, the size of the pupil changes in response to the amount of light hitting the retina (Campbell & Gregory, 1960). However, the pupil does not merely reflect accommodation. Instead, the size of the pupil is controlled by a balance between parasympathetic and sympathetic pathways that reflect both light-driven accommodation and central modulation. Several studies have shown that pupil size is associated with arousal (Joshi & Gold, 2019; Mathôt, 2018; C. A. Wang & Munoz, 2015). In the mammalian brain, arousal has been largely associated with the activity of the locus coeruleus (LC) (Aston-Jones & Cohen, 2005; Sara, 2009; van den Brink, Pfeffer, & Donner, 2019). This small structure in the pons contains a dense population of noradrenergic neurons and is the primary source of norepinephrine (NE) to the central nervous system. Recent neurophysiological work carried out in rodents and non-human primates has shown that pupil size is significantly associated with the spiking responses of LC neurons (Breton-Provencher & Sur, 2019; Joshi, Li, Kalwani, & Gold, 2016; Reimer et al., 2016; Varazzani, San-Galli, Gilardeau, & Bouret, 2015). Under conditions of heightened arousal, increases in LC activity and NE concentration are accompanied by increases in pupil size (Aston-Jones & Cohen, 2005; Sara, 2009; van den Brink et al., 2019).

Voluntary saccades occur ∼3 times per second to bring novel objects onto the high-resolution fovea (Kowler, 2011). The characteristics of these saccades, such as the reaction time to initiate the saccade, and the velocity reached during the saccade, have similarly been used to index global changes in arousal (Di Stasi et al., 2013). For example, when arousal is increased by delivering a startling auditory stimulus prior to the execution of a saccade, reaction time decreases and saccade velocity increases (Castellote, Kumru, Queralt, & Valls-Solé, 2007; Deuter, Schilling, Kuehl, Blumenthal, & Schachinger, 2013; DiGirolamo, Patel, & Blaukopf, 2016; Kristjánsson, Vandenbroucke, & Driver, 2004). Another metric that has been linked to global brain states, although to a lesser extent than reaction time and saccade velocity, is microsaccade rate. These small involuntary saccades are generated at a rate of 1-2Hz through the activity of neurons in the superior colliculus (SC) (Rolfs, 2009). Evidence suggests that microsaccade rate decreases with increased cognitive effort on a range of behavioral tasks (Gao, Yan, & Sun, 2015; Siegenthaler et al., 2014; Valsecchi, Betta, & Turatto, 2007; Valsecchi & Turatto, 2009). Taken together, these results suggest that the action of the eyes, be it when they are relatively stable and when they move, can be used to index global brain states.

A number of studies have related the activity of single neurons in many regions of the brain to pupil size (Joshi et al., 2016; Reimer et al., 2014), microsaccade rate (Bair & O’Keefe, 1998; Chen, Ignashchenkova, Thier, & Hafed, 2015; Herrington et al., 2009; Leopold & Logothetis, 1998; Lowet et al., 2018; Martinez-Conde, Macknik, & Hubel, 2000; Snodderly, Kagan, & Moshe, 2001), reaction time (Cook & Maunsell, 2002; Hanes & Schall, 1996; Khanna, Snyder, & Smith, 2019; Roitman & Shadlen, 2002; Steinmetz & Moore, 2019; Supèr & Lamme, 2007) and saccade velocity (Huang & Lisberger, 2009; O’Leary & Lisberger, 2012). However, if changes in these eye metrics are driven by a shift in an underlying internal state, then one might expect them all to be related to a common underlying neural activity pattern. Recent work in our laboratory used dimensionality reduction to identify a dominant mode of neural activity called slow drift that was related to behavior in a change detection task and present in both visual and prefrontal cortex in the macaque (Cowley et al., 2020). If such a prevalent change in brain-wide neural activity was truly reflective of a changing internal state, then it might also be reflective of wide-ranging changes in behavior. This motivated us to ask if our measure of the brain’s internal state (“slow drift”) is related to the activity of the eyes in different behavioral tasks. We recorded the spiking responses of populations of neurons in V4 while monkeys performed a change detection task and PFC while the same subjects performed a memory-guided saccade task. On both tasks, slow drift was associated with a pattern that is indicative of changes in the subjects’ arousal level over time. When pupil size increased, microsaccade rate and reaction time decreased while saccade velocity increased. These results show that the collective action of the eyes is associated with a dimension of neural activity that is pervasive and task-independent. They support the view that slow drift indexes a global brain state that manifests in the movement of the eyes and the size of the pupil.

## Results

To determine if observation of the eyes could provide insight into the internal state associated with slow drift, we recorded the spiking responses of populations of neurons in two macaque monkeys using 100-channel “Utah” arrays. We recorded from neurons in 1) V4 while the subjects performed an orientation-change detection task (Figure 1A); and 2) PFC while they performed a memory-guided saccade task (Figure 1B). Behavioral data for both subjects on the change detection task (Snyder, Yu, & Smith, 2018) and the memory-guided saccade task (Khanna, Scott, & Smith, 2019) has been published before in reports analyzing distinct aspects of the experiments described in this study. Here, the primary goal was to determine whether the neural population activity was related to eye metrics in a predictable manner across tasks. We analyzed data recorded in V4 on the change detection task because neural activity in midlevel visual areas has long been associated with performance on perceptual decision-making tasks (Shadlen, Britten, Newsome, & Movshon, 1996). Similarly, neural activity in PFC is correlated with performance on memory-guided saccade tasks (Funahashi, Bruce, & Goldman-Rakic, 1989). Four eye metrics were recorded during each session: pupil size, microsaccade rate, reaction time and saccade velocity. These metrics were chosen because they have been used extensively to index global changes in brain state (see Introduction), and can be measured easily and accurately with an infrared eye tracker (Kimmel et al., 2012). We made each of these four measurements on every trial of both behavioral tasks (Figure 1). Similar trends were observed in both subjects in a number of individual sessions (Figure 4 – figure supplement 2 and Figure 4 – figure supplement 3). For this reason, and to enhance statistical power, the data for Monkey 1 (20 sessions) and Monkey 2 (16 sessions) was combined for each task.

**Figure 1.**
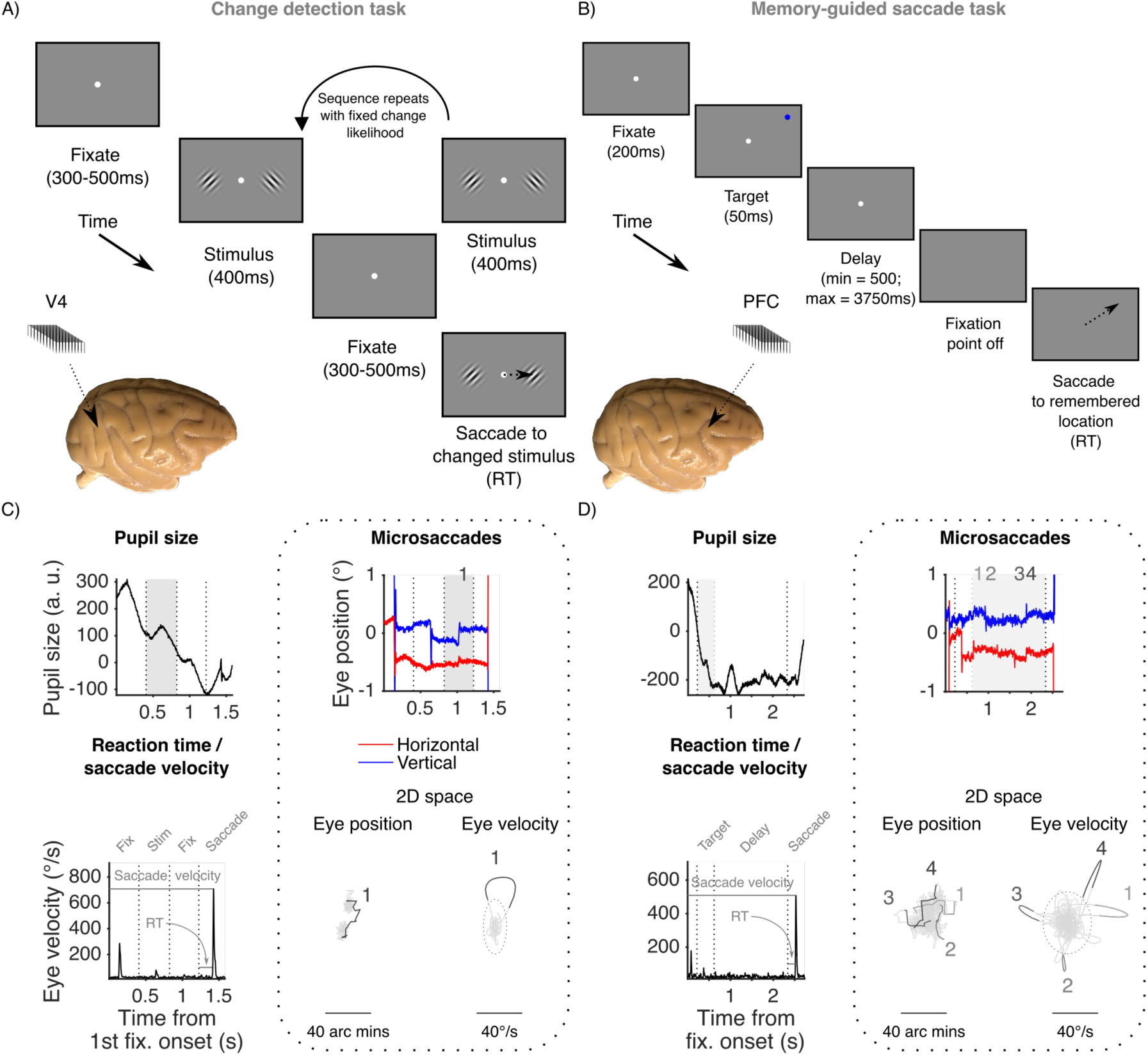
Experimental methods. (A) Change detection task. After an initial fixation period, a sequence of stimuli (orientated Gabors separated by fixation periods) was presented. The subjects’ task was to detect an orientation change in one of the stimuli and make a saccade to the changed stimulus. We recorded neural activity from V4 while the animals performed the change detection task using 100-channel “Utah” arrays (inset). (B) Memory-guided saccade task. After an initial fixation period, a target stimulus was presented at 1 of 40 locations followed by a delay period. The central point was then extinguished prompting the subjects to make a saccade to the remembered target location. Neural activity was recorded in PFC while the animals performed the memory-guided saccade task using Utah arrays (inset). (C) Measuring eye metrics on the change detection task. Mean pupil size was recorded during stimulus periods, whereas microsaccade rate was measured during periods of steady fixation (except for the initial fixation period, see *Methods*). Microsaccades (emboldened in two-dimensional eye position and eye velocity space) were defined as eye movements that exceeded a threshold (dashed circle) for 6ms (Engbert & Kliegl, 2003). Reaction time was the time taken to make a saccade to the changed stimulus. Saccade velocity was the peak velocity of the saccade. (D) Measuring eye metrics on the memory-guided saccade task. Mean pupil size was recorded during the presentation of the target stimulus, whereas microsaccade rate was measured during the delay period. Reaction time was the time taken to make a saccade to the remembered target location. Saccade velocity was the peak of velocity of the saccade. RT = reaction time. Figure supplement 1. Scatter plots and histograms showing the relationship between microsaccade amplitude and peak velocity.

### Correlations between the eye metrics over time

First, we investigated if the different measures of the eyes were themselves correlated over time during performance of the behavioral task. A large body of work has shown that arousal is associated with changes in the action of the eyes, be it when they are relatively stable or when they move. As described above, increases in arousal are typically accompanied by increases in pupil size and saccade velocity, and concomitant decreases in microsaccade rate and reaction time. Given that arousal is a domain-general phenomenon, one might expect a similar pattern to emerge on different behavioral tasks. To explore whether or not this was the case, we binned our eye data using a 30-minute sliding window stepped every 6 minutes (Figure 2A and Figure 2B). The width of the window, and the step size, were chosen to isolate slow changes over time based on previous research. They were the same as those used by Cowley et al. (2020), which meant direct comparisons could be made across studies. An example session from the same subject on the change detection task and the memory-guided saccade task is shown in Figure 2C and Figure 2D, respectively. In the example sessions shown across both tasks a characteristic pattern was observed that was indicative of slow changes in the subject’s arousal level over time. Specifically, a large tonic pupil size (measured during stimulus periods on the change detection task and target presentations on the memory-guided saccade task) was associated with high saccade velocity, shorter reaction times, and a lower rate of microsaccades.

**Figure 2.**
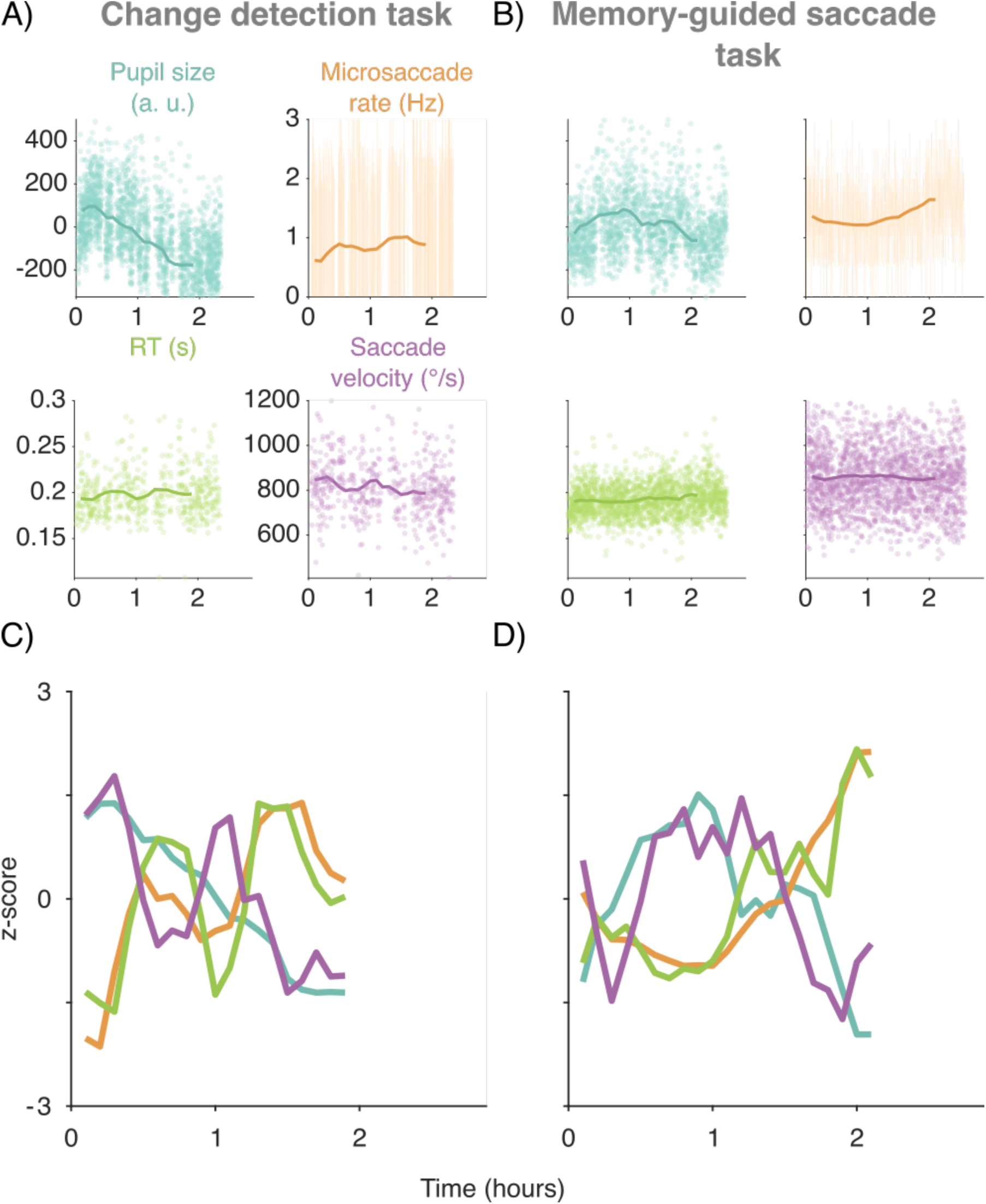
Isolating slow fluctuations in the eye metrics. (A) Change detection task. Each data point corresponds to a measurement from a single trial. To isolate slow changes, the data was binned using a 30-minute sliding window stepped every 6 minutes (solid line) (Cowley et al., 2020). (B) Same as (A) but for the memory-guided saccade task. Note in (B) and (C) that data points with a SD ∼3 times greater than the mean are not shown for illustration purposes. (C) Example session from Monkey 1 on the change detection task. Each metric has been z-scored for illustration purposes. (D) Example session from the same subject on the memory-guided saccade task.

Next, we explored if a similar pattern was found across all sessions and both animals. We computed correlations (Pearson product-moment correlation coefficient) between all combinations of the four eye metrics for each session, and compared that distribution of Pearson’s r values to shuffled distributions using permutation tests (two-sided, difference of medians). Consistent with the pattern of results observed in several individual sessions (Figure 3 – figure supplement 1 and Figure 3 – figure supplement 2), we found significant interactions among the four eye metrics. Pupil size was significantly and negatively correlated with microsaccade rate (median r = −0.46; p < 0.001) and reaction time (median r = −0.18; p = 0.020) on the change detection task (Figure 3A and Figure 3 – figure supplement 1) and memory-guided saccade task (Figure 3B and Figure 3 – figure supplement 2, median r = −0.56; p < 0.001 for microsaccade rate and median r = −0.15; p = 0.011 for reaction time). We did not observe significant across-session trends in the correlation between pupil size and saccade velocity (change detection task: r = 0.03, p = 0.987; memory-guided saccade task: r = 0.09, p = 0.523 in Figure 3A-B), although saccade velocity was itself correlated with reaction time in both tasks (change detection task: r = −0.52, p < 0.001; memory-guided saccade task: r = −0.61, p < 0.001). In addition, on the change detection task, we found a significant positive correlation between microsaccade rate and rection time (median r = 0.14; p = 0.004). Taken together, these results demonstrate that changes in the subjects’ arousal level were accompanied by changes in the action of the eyes. This was true of pupil size, microsaccade rate, reaction time, and to a lesser extent, saccade velocity, and suggests that these eye metrics may be used to index global changes in brain state. This motivated us to ask next whether there were neural correlates of these behavioral signatures.

**Figure 3.**
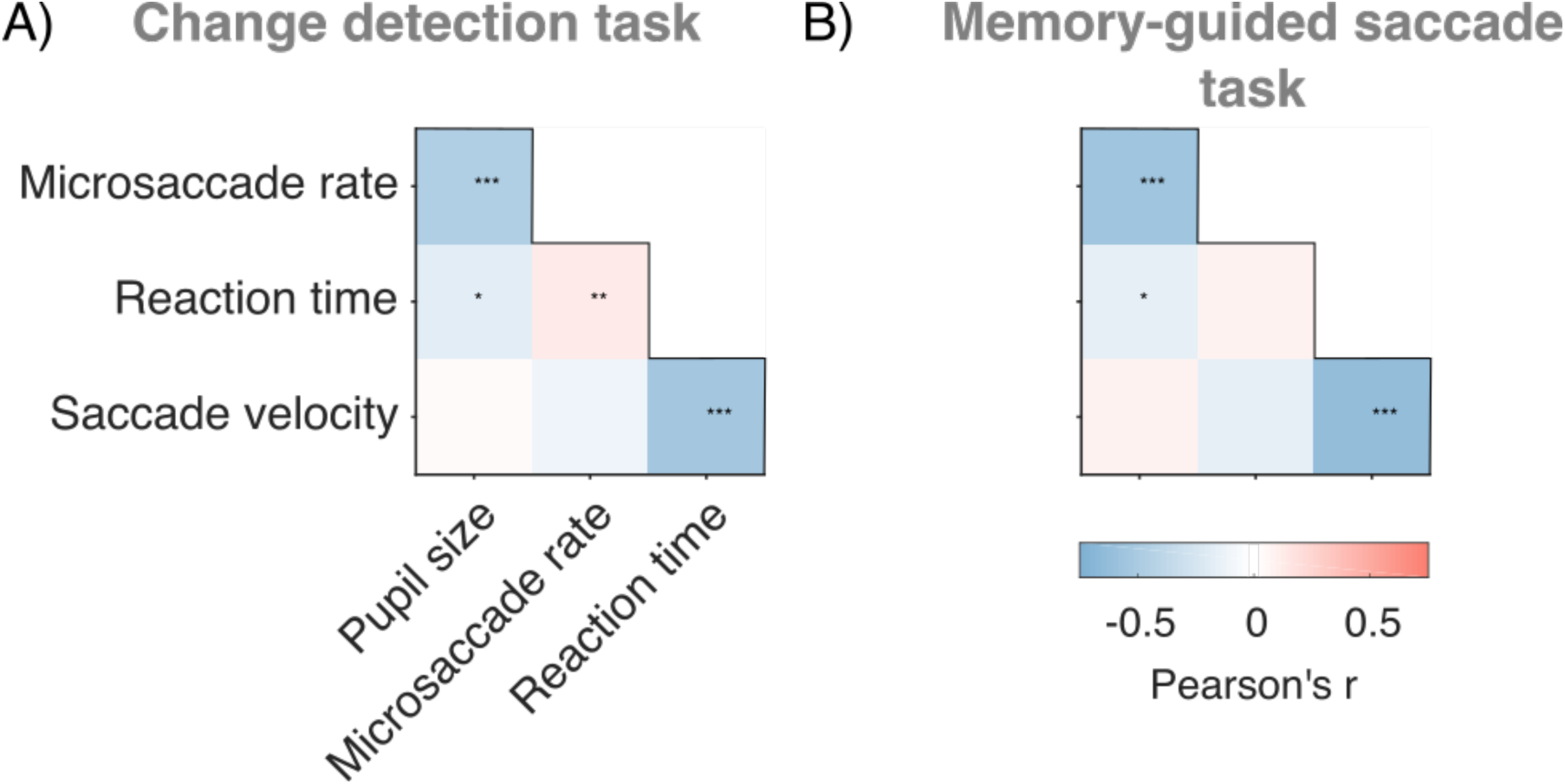
Correlations between the eye metrics over time. (A) Change detection task. Correlation matrix showing median r values across sessions between the four eye metrics. (B) Same as (A) but for the memory-guided saccade task. In (A) and (B) actual distributions of r values were compared to shuffled distributions using two-sided permutation tests (difference of medians). p < 0.05*, p < 0.01**, p < 0.001***. Figure supplement 1. Three example sessions from Monkey 1 and histograms showing actual and shuffled distributions of r values across sessions on the change detection task. Figure supplement 2. Three example sessions from Monkey 1 and histograms showing actual and shuffled distributions of r values across sessions on the memory-guided saccade task.

### Correlation between the eye metrics and slow drift over time

Previously we reported a slow fluctuation in neural activity in V4 and PFC that we termed ‘slow drift’ (Cowley et al., 2020). We found that this neural signature was related to the subject’s tendency to make impulsive decisions in a change detection task, ignoring sensory evidence (false alarms). Here, we wanted to investigate if the constellation of eye metrics we observed were associated with the neural signature of internal state that we termed ‘slow drift.’ To calculate slow drift, we binned spike counts in V4 (change detection task) and PFC (memory-guided saccade task) using the same 30-minute sliding window that had been used to bin the eye metric data (Figure 4A-B, see *Methods*). We then applied principal component analysis (PCA) to the data and estimated slow drift by projecting the binned residual spike counts along the first principal component (i.e., the loading vector that explained the most variance in the data). Because the sign of the loadings in PCA is arbitrary (Jollife & Cadima, 2016), the correlation between slow drift and a given eye metric in any session was equally likely to be positive or negative. This was problematic since we were interested in whether slow drift was associated with a characteristic pattern that is indicative of changes in the subjects’ arousal level over time i.e., increased pupil size and saccade velocity, and decreased microsaccade rate and reaction time. In order to establish consistency across sessions in the sign of the correlations, we constrained the slow drift in each session to have the same relationship to the spontaneous (change detection task = fixation periods; memory-guided saccade task = delay periods) and evoked (change detection task = stimulus periods; memory-guided saccade task = target presentations) activity of the neurons (Figure 4 – figure supplement 1, and see *Methods*). This served to align slow drift across sessions such that an increase in the slow drift value was associated with neural activity closer to the evoked pattern of response (i.e., typically higher firing rates).

**Figure 4.**
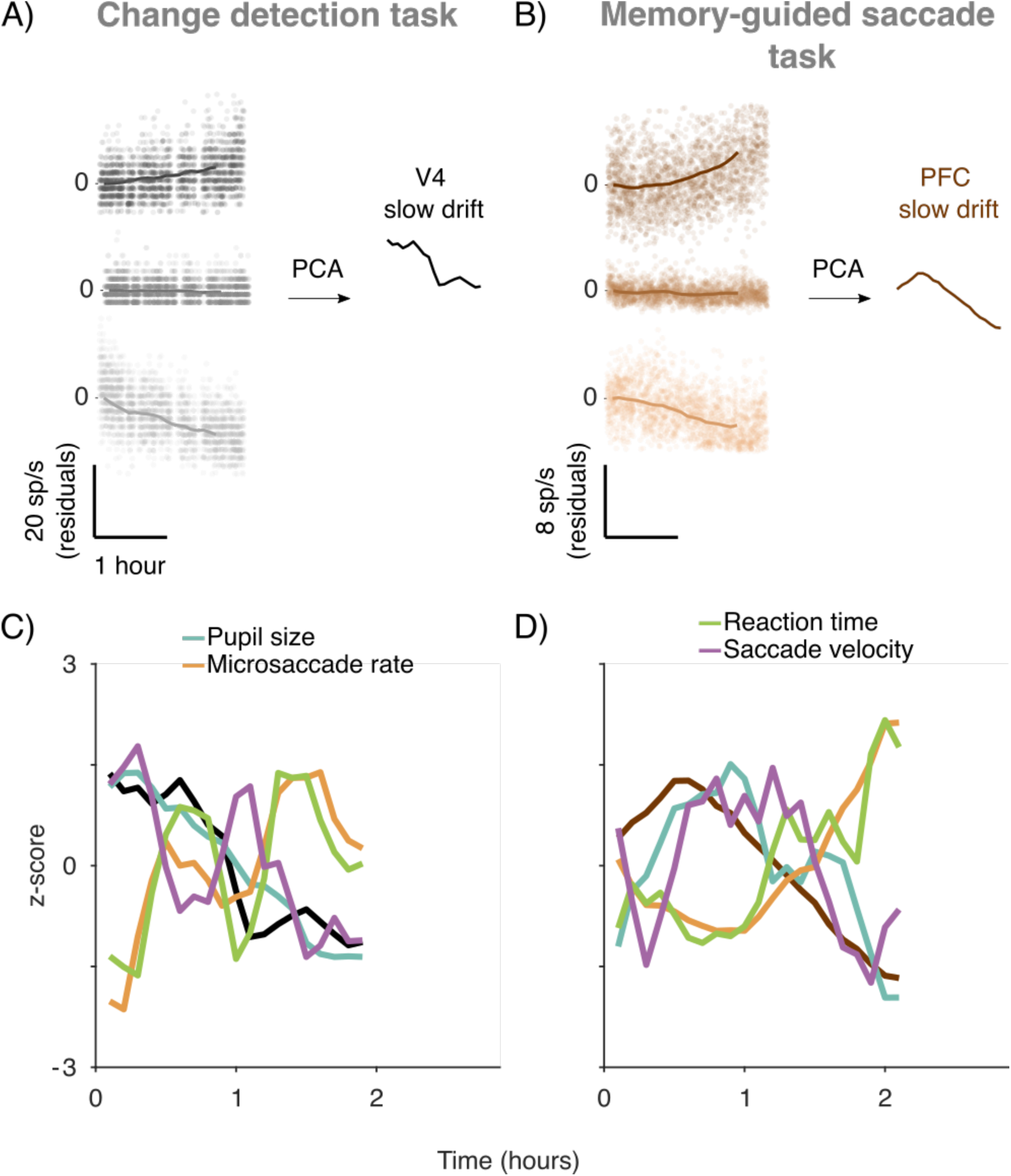
Calculating slow drift. (A) Change detection task. Three example neurons from a single session (Monkey 1). Each point represents the mean residual spike count during a 400ms stimulus period. The data was then binned using a 30-minute sliding window stepped every six minutes (solid line) so that direct comparisons could be made with the eye metrics. PCA was used to reduce the dimensionality of the data and slow drift was calculated by projecting binned residual spike counts along the first principal component. (B) Same as (A) but for the memory-guided saccade task (same subject). (C) Example session from Monkey 1 on the change detection task. Each metric has been z-scored for illustration purposes. (D) Example session from the same subject on the memory-guided saccade task. Figure supplement 1. Plots showing how slow drift was aligned across sessions. Figure supplement 2. Example sessions from Monkey 1 on the change detection task and the memory-guided saccade task. Figure supplement 3. Example sessions from Monkey 2 on the change detection task and the memory-guided saccade task. Figure supplement 4. Percent waveform variance of four example neurons recorded from Monkey 1 during a single session on the change detection task and the memory-guided saccade task.

We computed the slow drift of the neuronal population in each session using this method, and then compared it to the four eye measures. An example session is shown in Figure 4C-D for the same subject on the change detection task and the memory-guided saccade task, respectively (same sessions as in Figure 2). On both tasks, we found a characteristic pattern in which slow drift was positively associated with pupil diameter and saccade velocity, and negatively associated with microsaccade rate and reaction time.

Next, we explored if a similar pattern was found across sessions. We computed correlations (Pearson product-moment correlation coefficient) between slow drift and the eye metrics. Actual distributions of r values were then compared to shuffled distributions using permutation tests (two-sided, difference of medians). Because the slow drift was aligned across sessions based on the neural activity alone and not the behavior, the shuffled distributions are centered on a correlation value of zero. Consistent with the pattern of results observed in several individual sessions (Figure 4 – figure supplement 2 and Figure 4 – figure supplement 3), we found that slow drift in V4 on the change detection task (Figure 5A) was positively correlated with pupil diameter (median r = 0.46, p < 0.001) and saccade velocity (median r = 0.19, p = 0.022), and negatively correlated with microsaccade rate (median r = −0.28, p = 0.010). While no significant correlation was found between slow drift and reaction time in the data pooled across subjects (median r = −0.07; p = 0.572), one subject (Monkey 1) did exhibit a negatively correlation with reaction time (median r = −0.19, p = 0.039). An identical pattern of results was found on the memory-guided saccade task (Figure 5B), where slow drift in PFC was positively correlated with pupil diameter (median r = 0.20; p = 0.029) and saccade velocity (median r = 0.17, p = 0.042), and negatively correlated with microsaccade rate (median r = −0.42; p < 0.001) and reaction time (median r = −0.27; p = 0.008). These results show that pupil size, microsaccade rate, reaction time and saccade velocity are associated with slow drift across tasks. Thus, the slow drift was associated with a pattern that is indicative of changes in the subject’s arousal levels over time.

**Figure 5.**
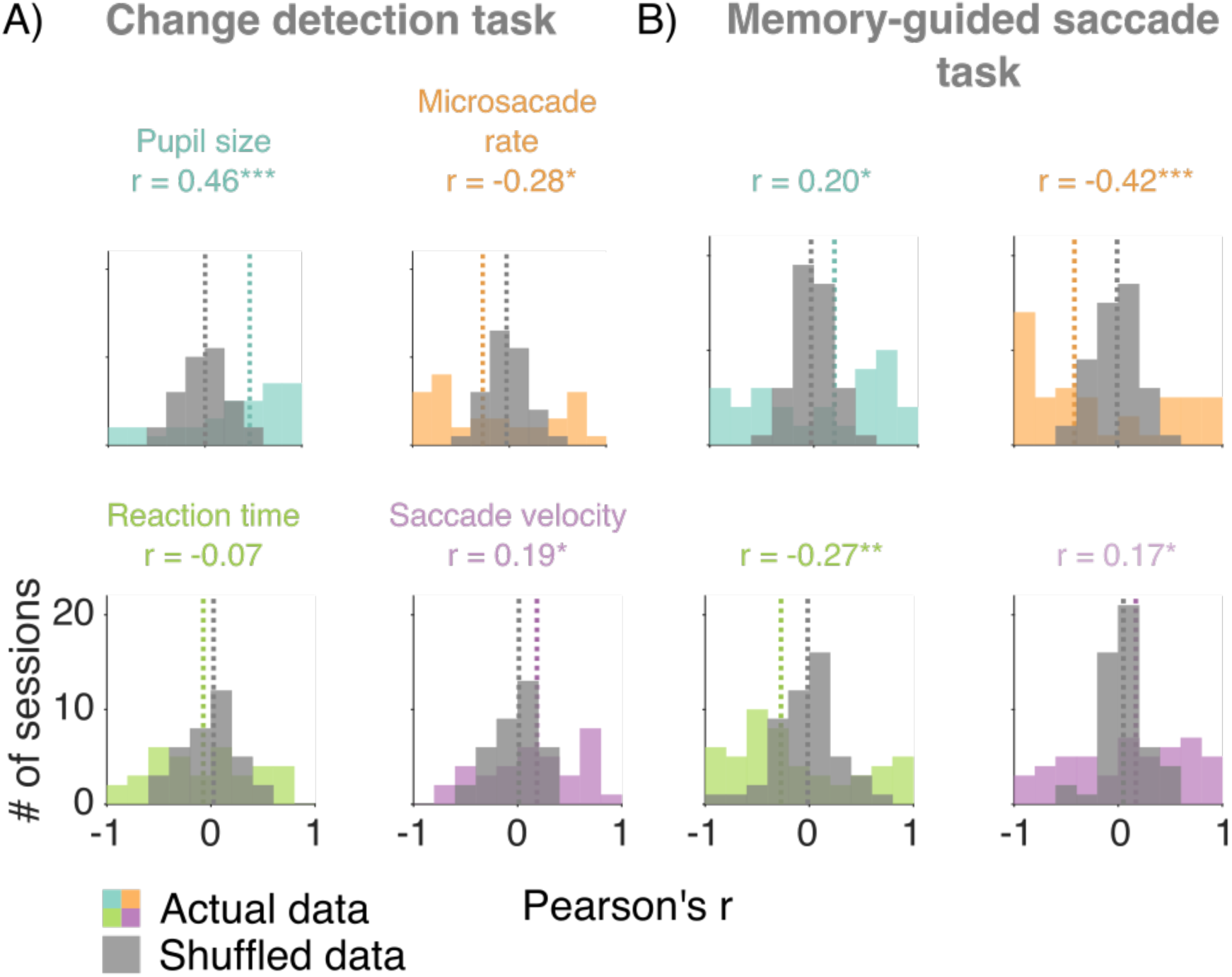
Correlations between the eye metrics and slow drift over time. (A) Change detection task. Histograms showing actual and shuffled distributions of r values. (B) Same as (A) but for the memory-guided saccade task. In (A) and (B) median r values across sessions are indicated by dashed lines (colored lines = actual data; gray lines = shuffled data). Actual distributions of r values were compared to shuffled distributions using two-sided permutation tests (difference of medians). p < 0.05*, p < 0.01**, p < 0.001***. Figure supplement 1. Histograms showing actual and shuffled distributions of r values computed to explore if simultaneously recorded PFC data was associated with the eye metrics on the change detection task.

## Discussion

In this study, we investigated if the size of the pupil and the movement of the eyes could be taken as an external signature of an internal brain state, a low-dimensional neural activity pattern called slow drift (Cowley et al., 2020). There is strong evidence that internal brain states such as slow drift can be measured in the population spiking activity of neurons, and that measurements of the eyes can provide important context into the behavior of subjects on perceptual and decision-making tasks. Hence, we were keen to determine whether we could directly link a neural measure of internal brain state acquired from the spiking activity of a population of neurons with external features of behavior. On two types of perceptual tasks, we found that slow drift was significantly correlated with a pattern of eye metrics that was indicative of changes in the subjects’ arousal level over time. Our results show that the action of the eyes, be it when they are relatively stable or when they move, is associated with a latent dimension of neural activity that is pervasive and task-independent.

As described above, decades of research have shown that eye metrics are related to task performance in a variety of contexts (Di Stasi et al., 2013; Joshi & Gold, 2019; Mathôt, 2018; C. A. Wang & Munoz, 2015). Heightened levels of arousal have been associated with increased pupil size and saccade velocity as well as decreased reaction time and microsaccade rate (Castellote et al., 2007; Deuter et al., 2013; DiGirolamo et al., 2016; Gao et al., 2015; Joshi et al., 2016; Siegenthaler et al., 2014; Valsecchi et al., 2007; Valsecchi & Turatto, 2009). Given that arousal is a global phenomenon, one might expect a common pattern across multiple behavioral tasks. In the present study, we addressed this issue by investigating the relationships between eye metrics on tasks designed to probe the mechanisms underlying perceptual decision-making (change detection task) and working memory (memory-guided saccade task). Results showed that pupil size was negatively correlated with microsaccade rate and reaction time on both tasks. Although no significant correlation was found between pupil size and saccade velocity, the overall pattern of results suggests that each subject’s arousal level was changing over time in a task-independent manner. These findings support the view that measuring properties related to the eyes can provide a non-invasive index of global brain states (Di Stasi et al., 2013; Joshi & Gold, 2019; Mathôt, 2018; C. A. Wang & Munoz, 2015). This motivated us to ask if they are also associated with slow drift.

Most studies that have explored the relationship between eye metrics and neural activity have used single-neuron (spike count, Fano factor) and pairwise statistics (r_sc_). However, numerous recent studies have shown that rich insight about cognitive processes (e.g., learning, decision-making, working memory, time perception) can be gained from analysis of the simultaneous activity of populations of neurons (Harvey, Coen, & Tank, 2012; Mante, Sussillo, Shenoy, & Newsome, 2013; Murray et al., 2017; Remington, Narain, Hosseini, & Jazayeri, 2018; Sadtler et al., 2014). In addition, recent work has shown that a low-dimensional representation of neural activity in the mouse can be used to index global changes in brain state. Stringer et al. (2019) applied PCA to data recorded from more than 10,000 neurons and found that fluctuations in the first principal component were significantly associated with whisking, pupil size, and running speed. In this study, we investigated if slow drift, a dominant mode of neural activity in macaque cortex is associated with the action of the eyes. On both tasks, we found that it was correlated with a pattern of eye metrics that is indicative of changes in the subject’s arousal level over time. Our results, coupled with those of Stringer et al. (2019), suggest that much can be learned about global brain states, as well as cognitive processes, when high-dimensional population activity is reduced to a low-dimensional subspace (Cunningham & Yu, 2014). A key question for future research is whether latent dimensions of neural activity in the cortex are associated with activity in subcortical brain regions? One might expect this to be the case given that slow drift was significantly correlated with pupil size on both tasks.

Evidence suggests that pupil size is associated with activity in the LC, a subcortical structure that regulates arousal by releasing NE in a diverse manner throughout much of the brain (Aston-Jones & Cohen, 2005; Sara, 2009; van den Brink et al., 2019). That slow drift can be used to index global brain states suggests that it might be associated with LC activity. Although the LC is limited in size (∼3mm rostrocaudally) and buried deep within the brainstem (German & Bowden, 1975; Sharma et al., 2010), several studies have successfully recorded from single neurons in LC of the macaque (Aston-Jones, Rajkowski, Kubiak, & Alexinsky, 1994; Clayton, Rajkowski, Cohen, & Aston-Jones, 2004; Joshi et al., 2016; Kalwani, Joshi, & Gold, 2014; Varazzani et al., 2015). In addition, LC can be activated using optogenetics (Carter et al., 2010; Hayat et al., 2020; Li et al., 2016; Quinlan et al., 2019), electrical microstimulation (Joshi et al., 2016; Liu, Rodenkirch, Moskowitz, Schriver, & Wang, 2017; Reimer et al., 2016), and pharmacological manipulations (Liu et al., 2017; Vazey & Aston-Jones, 2014). Thus, it should be possible to alter the course of slow drift in the cortex using some, if not all, of these methods. As well as having a significant effect on pupil size, our results predict that activating the LC, directly or indirectly, may lead to task-independent changes in microsaccade rate, reaction time and saccade velocity.

Evidence suggests that microsaccades, reaction time and saccade velocity are associated with neural activity in the SC (Gandhi & Katnani, 2011). This layered structure, located at the roof of the brain stem, plays a critical role in transforming sensory information into eye movement commands. Population recordings have been successfully performed in the SC using linear probes (Massot, Jagadisan, & Gandhi, 2019), and it thus might be possible to identify dominant patterns of neural activity in SC using dimensionality reduction (Cunningham & Yu, 2014). Our results predict that slow drift in the cortex should be significantly correlated with slow drift in the SC. However, this might depend upon the mixture of SC neurons in the recorded population. We previously suggested that slow drift must be removed at some stage before motor commands are issued to prevent unwanted eye movements (Cowley et al., 2020). Thus, one might not expect a correlation between slow drift in the cortex and slow drift in deep-layer SC neurons that fire vigorously prior to the execution of a saccade, and relay motor commands to downstream nuclei innervating the oculomotor muscles (Sparks & Hartwich-Young, 1989). An alternative possibility is that slow drift is not removed, but instead occupies an orthogonal subspace in the SC that is not read out by downstream nuclei. This is not beyond the realm of possibility given that an identical scheme appears to exist in the skeletomotor system to stop preparatory signals reaching the muscles (Ames & Churchland, 2019; Elsayed, Lara, Kaufman, Churchland, & Cunningham, 2016; Kaufman, Churchland, Ryu, & Shenoy, 2014; Stavisky, Kao, Ryu, & Shenoy, 2017). Further research is needed to disentangle these possibilities.

Studies in the fields of psychology and neuroscience have mainly used pupil size to index arousal, but metrics such as heart rate (HR) and galvanic skin response (GSR) are also associated with global brain states. For example, Wang et al. (2018) measured pupil size, HR and GSR while human subjects viewed emotional face stimuli specifically designed to evoke fluctuations in arousal. Results showed that all three metrics were positively correlated prior to the presentation of the stimuli. That is, when pupil size was large, and the subjects were in heightened state of arousal, HR and GSR increased. In addition, it has been suggested that pre-stimulus oscillations in the alpha band can be used to index global brain states as they are inversely related to performance on visual detection tasks. Several studies using electroencephalography (EEG) have shown that the probability of detecting near-threshold stimuli increases when pre-stimulus power in the alpha band is low (Benwell et al., 2017; Samaha, Iemi, & Postle, 2017; Van Dijk, Schoffelen, Oostenveld, & Jensen, 2008). Simultaneous recordings of spiking activity and EEG in awake behaving monkeys are rare. However, our results predict that slow drift should be associated with pre-stimulus alpha oscillations.

Research has also uncovered a link between alpha oscillations and microsaccades (Bellet, Chen, & Hafed, 2017). This effect has been attributed to changes in spatial attention, which have a profound effect on microsaccade direction. For example, Lowet et al. (2018) found that attention-related modulation of spiking responses, Fano factor and r_sc_ only occur following a microsaccade in the direction of an attended stimulus. In the present study, we found that slow drift was significantly correlated with microsaccade rate on both tasks. This is unlikely to be explained by changes in spatial attention as both Cowley et al. (2020) and Rabinowitz et al. (2015) found that slow drift on a change detection task was not associated with the blocks of trials used to cue spatial attention inside and outside the RF. In addition, the correlation between slow drift and microsaccade rate in the present study was lower on the change detection task (r = −0.27) than the memory-guided saccade task (r = −0.37). One would not have expected this to be the case if slow drift was mediated by changes in spatial attention. These results raise the possibility that spatial attention and arousal have differential effects on microsaccades. Attention might specifically affect microsaccade direction, whereas arousal might affect microsaccade rate irrespective of direction. Further research is needed to test this hypothesis.

In summary, we investigated if properties related to the eyes are associated with slow drift: a low-dimensional pattern of neural activity that was recently identified in the macaque cortex by Cowley et al. (2020). On both tasks, we found that slow drift was significantly associated with a pattern of eye metrics that is indicative of changes in the subjects’ arousal level over time. These results demonstrate that the collective action of the eyes is associated with a latent dimension of neural activity that is pervasive and task-independent. They suggest that slow drift can be used to index global changes in brain state over time. Further research is necessary to determine the origins of this slow drift in population activity. A key question for future work will be to determine the mechanisms by which slow drift influences behavior in some instances (e.g., when arousal level drives an urgent response) and is circumvented in others (e.g., when an accurate perceptual judgement must be made regardless of arousal level).

## Methods

### Subjects

Two adult rhesus macaque monkeys (*Macaca mulatta*) were used in this study. Surgical procedures to chronically implant a titanium head post (to immobilize the subjects’ head during experiments) and microelectrode arrays were conducted in aseptic conditions under isoflurane anesthesia, as described in detail by Smith and Sommer (2013). Opiate analgesics were used to minimize pain and discomfort during the perioperative period. Neural activity was recorded using 100-channel “Utah” arrays (Blackrock Microsystems) in V4 (Monkey 1 = right hemisphere; Monkey 2 = left hemisphere) and PFC (Monkey 1 = right hemisphere; Monkey 2 = left hemisphere) while the subjects performed the change detection task (Figure 1A). Note that this is the same dataset used by Snyder et al. (2018) and Cowley et al. (2020). On the memory-guided saccade task (Figure 1B), neural activity was only recorded in PFC (Monkey 1 = left hemisphere; Monkey 2 = left hemisphere). Note that the data presented here are a superset of the data presented in Khanna et al. (2019). The only difference between the memory-guided saccade data presented here and that previous study is that here we also analyzed neural activity from additional sessions in Monkey 1 after a new array was implanted in left PFC. The sessions were also longer, and particularly well suited to analyze slow fluctuations in neural activity. The arrays comprised a 10×10 grid of silicon microelectrodes (1 mm in length) spaced 400 μm apart. Experimental procedures were approved by the Institutional Animal Care and Use Committee of the University of Pittsburgh and were performed in accordance with the United States National Research Council’s Guide for the Care and Use of Laboratory Animals.

### Microelectrode array recordings

Signals from each microelectrode in the array were amplified and band-pass filtered (0.3–7500 Hz) by a Grapevine system (Ripple). Waveform segments crossing a threshold (set as a multiple of the root mean square noise on each channel) were digitized (30KHz) and stored for offline analysis and sorting. First, waveforms were automatically sorted using a competitive mixture decomposition method (Shoham, Fellows, & Normann, 2003). They were then manually refined using custom time amplitude window discrimination software ((Kelly et al., 2007), code available at https://github.com/smithlabvision/spikesort), which takes into account metrics including (but not limited to) waveform shape and the distribution of interspike intervals. A mixture of single and multiunit activity was recorded, but we refer here to all units as “neurons”. On the change detection task, the mean number of V4 neurons across sessions was 70 (SD = 11) for Monkey 1 and 31 (SD = 16) for Monkey 2, whereas the mean number of PFC neurons across sessions was 84 (SD = 15) for Monkey 1 and 90 (SD = 20) for Monkey 2. On the memory-guided saccade task, the mean number of PFC neurons across sessions was 54 (SD = 11) for Monkey 1 and 38 (SD = 19) for Monkey 2.

### Visual stimuli

Visual stimuli were generated using a combination of custom software written in MATLAB (The MathWorks) and Psychophysics Toolbox extensions (Brainard, 1997; Pelli, 1997; Kleiner et al., 2007). They were displayed on a CRT monitor (resolution = 1024 X 768 pixels; refresh rate = 100Hz), which was viewed at a distance of 36cm and gamma-corrected to linearize the relationship between input voltage and output luminance using a photometer and look-up-tables.

## Behavioral tasks

### Orientation-change detection task

Subjects fixated a central point (diameter = 0.6°) on the monitor to initiate a trial (Figure 1A). Each trial comprised a sequence of stimulus periods (400ms) separated by fixation periods (duration drawn at random from a uniform distribution spanning 300-500ms). The 400ms stimulus periods comprised pairs of drifting full-contrast Gabor stimuli. One stimulus was presented in the aggregate receptive field (RF) of the recorded V4 neurons, whereas the other stimulus was presented in the mirror-symmetric location in the opposite hemifield. Although the spatial (Monkey 1 = 0.85cycles/°; Monkey 2 = 0.85cycles/°) and temporal frequencies (Monkey 1 = 8cycles/s; Monkey 2 = 7cycles/s) of the stimuli were not optimized for each individual V4 neuron they did evoke a strong response from the population. The orientation of the stimulus in the aggregate RF was chosen at random to be 45 or 135°, and the stimulus in the opposite hemifield was assigned the other orientation. There was a fixed probability (Monkey 1 = 30%; Monkey 2 = 40%) that one of the Gabors would change orientation by ±1, ±3, ±6, or ±15° on each stimulus presentation. The sequence continued until the subject) made a saccade to the changed stimulus within 700ms (“hit”); 2) made a saccade to an unchanged stimulus (“false alarm”); or 3) remained fixating for more than 700ms after a change occurred (“miss”). If the subject correctly detected an orientation change, they received a liquid reward. In contrast, a time-out occurred if the subject made a saccade to an unchanged stimulus delaying the beginning of the next trial by 1s. It is important to note that the effects of spatial attention were also investigated (although not analyzed in this study) by cueing blocks of trials such that the orientation change was 90% more likely to occur within the aggregate V4 RF than the opposite hemifield.

### Memory-guided saccade task

Subjects fixated a central point (diameter = 0.6°) on the monitor to initiate a trial (Figure 1B). After fixating within a circular window (diameter = 2.4° and 1.8° for Monkey 1 and Monkey 2, respectively) for 200ms, a target stimulus (diameter = 0.8°) was presented for 50ms (except for 1 session in which it was presented for 400ms). The target stimulus appeared at 1 of 8 angles separated by 45°, and 1 of 5 eccentricities, yielding 40 conditions in total. After the target stimulus had been presented, subjects were required to maintain fixation for a delay period. For Monkey 1, the duration of the delay period was either 1) drawn at random from a distribution spanning 1200-3750ms; or 2) fixed at 1400 or 2000ms. For Monkey 2, the duration of the delay period was 500ms. If steady fixation was maintained throughout the delay period, the central point was extinguished prompting the subjects to make a saccade to the remembered target location. The subjects had 500ms to initiate a saccade. Once the saccade had been initiated, they had a further 200ms to reach the remembered target location. To receive a liquid reward, the subjects’ gaze had to be maintained within a circular window centered on the target location (diameter = 4 and 2.7° for Monkey 1 and Monkey 2, respectively) for 150 ms. In a subset of sessions, the target was briefly reilluminated, after the fixation point was extinguished and the saccade had been initiated, to aid in saccade completion.

### Eye tracking

Eye position and pupil diameter were recorded monocularly at a rate of 1000Hz using an infrared eye tracker (EyeLink 1000, SR Research).

### Microsaccade detection

Microsaccades were defined as eye movements that exceeded a velocity threshold of 6 times the standard deviation of the median velocity for at least 6ms (Engbert & Kliegl, 2003). They were required to be separated in time by at least 100ms. In addition, we removed microsaccades with an amplitude greater than 1° and a velocity greater than 100°/s. To assess the validity of our microsaccade detection method, the correlation (Pearson product-moment correlation coefficient) between the amplitude and peak velocity of detected microsaccades (i.e., the main sequence) was computed for each session (Figure 1 – figure supplement 1). The mean correlation between microsaccade amplitude and peak microsaccade velocity across sessions was 0.86 (SD = 0.07) for the change detection task and 0.83 (SD = 0.04) for the memory-guide saccade task. These findings indicate that our microsaccade detection algorithm was robust (Zuber, Stark, & Cook, 1965).

## Eye metrics

### Change detection task

Mean pupil diameter was measured during stimulus periods, whereas microsaccade rate was measured during fixation periods (Figure 1C). We did not include the first fixation period when measuring microsaccade rate in the change-detection task. As can be seen in Figure 1C, there was an increase in eye position variability during this period resulting from fixation having been established a short time earlier (300-500ms). Such variability was not present in proceeding fixation periods. Reaction time and saccade velocity were measured on trials in which the subjects were rewarded for correctly detected an orientation change. Reaction time was defined as the time from when the change occurred to the time at which the saccade exceeded a velocity threshold of 100°/s. Saccade velocity was the peak velocity of the saccade to the changed stimulus. To isolate slow changes in the eye metrics over time the data for each session was binned using a 30-minute sliding window stepped every 6 minutes (Figure 2A).

### Memory-guided saccade task

Pupil size, microsaccade rate, reaction time and saccade velocity were measured on trials in which the subjects received a liquid reward for making a correct saccade to the remembered target location. Mean pupil diameter was measured during the presentation of the target stimulus, whereas microsaccade rate was measured during the delay period (Figure 1D). Reaction time was defined as the time from when the fixation point was extinguished to the time at which the saccade reached a threshold of 100 °/s. Saccade velocity was the peak of velocity of the saccade to the remembered target location. As in the change-detection task, the data for each session was binned using a 30-minute sliding window stepped every 6 minutes (Figure 2B).

## Calculating slow drift

### Orientation-change detection task

The spiking responses of populations of neurons in V4 were measured during a 400ms period that began 50ms after stimulus presentation (Figure 4A). To control for the fact that some neurons had a preference for one orientation (45 or 135°) over the other residual spike counts were calculated. We subtracted the mean response for a given orientation across the entire session from individual responses to that orientation. To isolate slow changes in neural activity over time, residual spike counts for each V4 neuron were binned using a 30-minute sliding window stepped every 6 minutes (Cowley et al., 2020). PCA was then performed to reduce the high-dimensional residual data to a smaller number of variables (Cunningham & Yu, 2014). Slow drift in V4 was estimated by projecting the binned residual spike counts for each neuron along the first principal component (Cowley et al., 2020). As described above, the spiking responses of neurons in PFC were simultaneously recorded on the change detection task. When PFC slow drift was calculated using the method described above, we found it to be significantly associated with V4 slow drift (median r = 0.95, p < 0.001), consistent with previous results (Cowley et al., 2020). On the change detection task, we investigated the relationships between the eye metrics and V4 slow drift. However, a very similar pattern of results was found when slow drift was calculated using simultaneously recorded PFC data (Figure 5 – figure supplement 1).

### Memory-guided saccade task

The spiking responses of populations of neurons in PFC were measured during the delay period (Figure 4B). To control for the fact that some neurons had a preference for one target location over another, residual spike counts were calculated. We subtracted the mean response to a given target location across the entire session from individual responses to that location. To isolate slow changes in neural activity over time residual spike counts for each PFC neuron were binned using a 30-minute sliding window stepped every 6 minutes. PCA was then performed, and slow drift was estimated by projecting the binned residual spike counts for each neuron along the first principal component.

### Controlling for neural recording instabilities

To rule out the possibility that slow drift arose due to recording instabilities (e.g., the distance between the neuron and the microelectrodes changing slowly over time) we only included neurons with stable waveform shapes throughout a session. This was quantified by calculating percent waveform variance for each neuron (Figure 4 – figure supplement 4). First, the session was divided into 10 non-overlapping time bins. A residual waveform was then computed for each time bin by subtracting the mean waveform across time bins. The variance of each residual waveform was divided by the variance of the mean waveform across time bins yielding 10 values (one of reach time bin). Percent waveform variance was defined as the maximum value across time bins. Neurons with a percent waveform variance greater than 10% were deemed as having unstable waveform shapes throughout a session. They were excluded from all analyses, consistent with previous research (Cowley et al., 2020).

### Aligning slow drift across sessions

As described above, slow drift was calculated by projecting binned residual spike counts along the first principal component (Cowley et al., 2020). The weights in a PCA can be positive or negative (Jollife & Cadima, 2016), which meant the sign of the correlation between slow drift and a given eye metric was arbitrary. Preserving the sign of the correlations was particularly important in this study because we were interested in whether slow drift was associated with a pattern that is indicative of changes in the subjects’ arousal levels over time i.e., increased pupil size and saccade velocity, and decreased microsaccade rate and reaction time. Thus, we had to devise a method to align slow drift across sessions. As expected, the mean evoked population response calculated during stimulus periods on the change detection task was significantly higher than the mean spontaneous population response calculated during fixation periods (Figure 4 – figure supplement 1). Similarly, on the memory-guided saccade task, the mean evoked population response calculated during target presentations was significantly higher than the mean spontaneous population response calculated during delay periods. To align slow drift for each individual session, we projected evoked and spontaneous responses onto the first principal component. The sign of the slow drift was flipped if the mean projection value for the evoked responses was less than that for the spontaneous responses – i.e., if the relationship described above for unprojected data did not hold true.

---

**Figure 1 – figure supplement 1.**
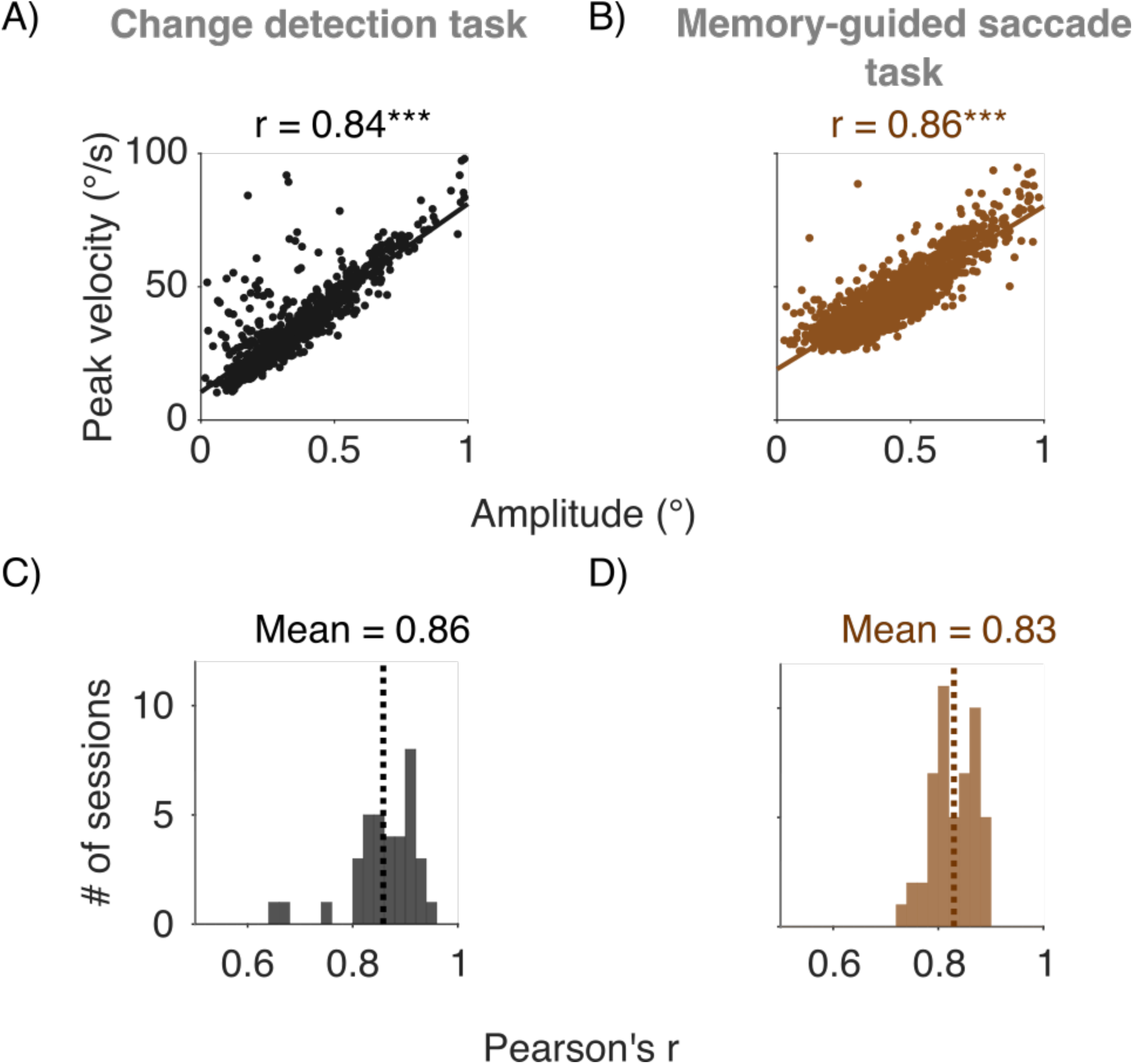
Assessing the validity of our microsaccade detection algorithm. The correlation between the amplitude and peak velocity of detected microsaccades was computed for each session. (A) Change detection task. Scatter plot showing the relationship between microsaccade amplitude and peak velocity for an example session on the change detection task. (B) Same as (A) but for the memory-guided saccade task (same subject). (C) Histogram showing the distribution of r values for the change detection task. (D) Same as (C) but for the memory-guided saccade task. In (C) and (D) dashed lines indicate the mean r value across sessions. On both tasks, the mean correlation between microsaccade amplitude and peak velocity was strong, indicating that our method of detecting microsaccades was robust (Zuber et al., 1965). p < 0.05*, p < 0.01**, p < 0.001***.

**Figure 3 - figure supplement 1.**
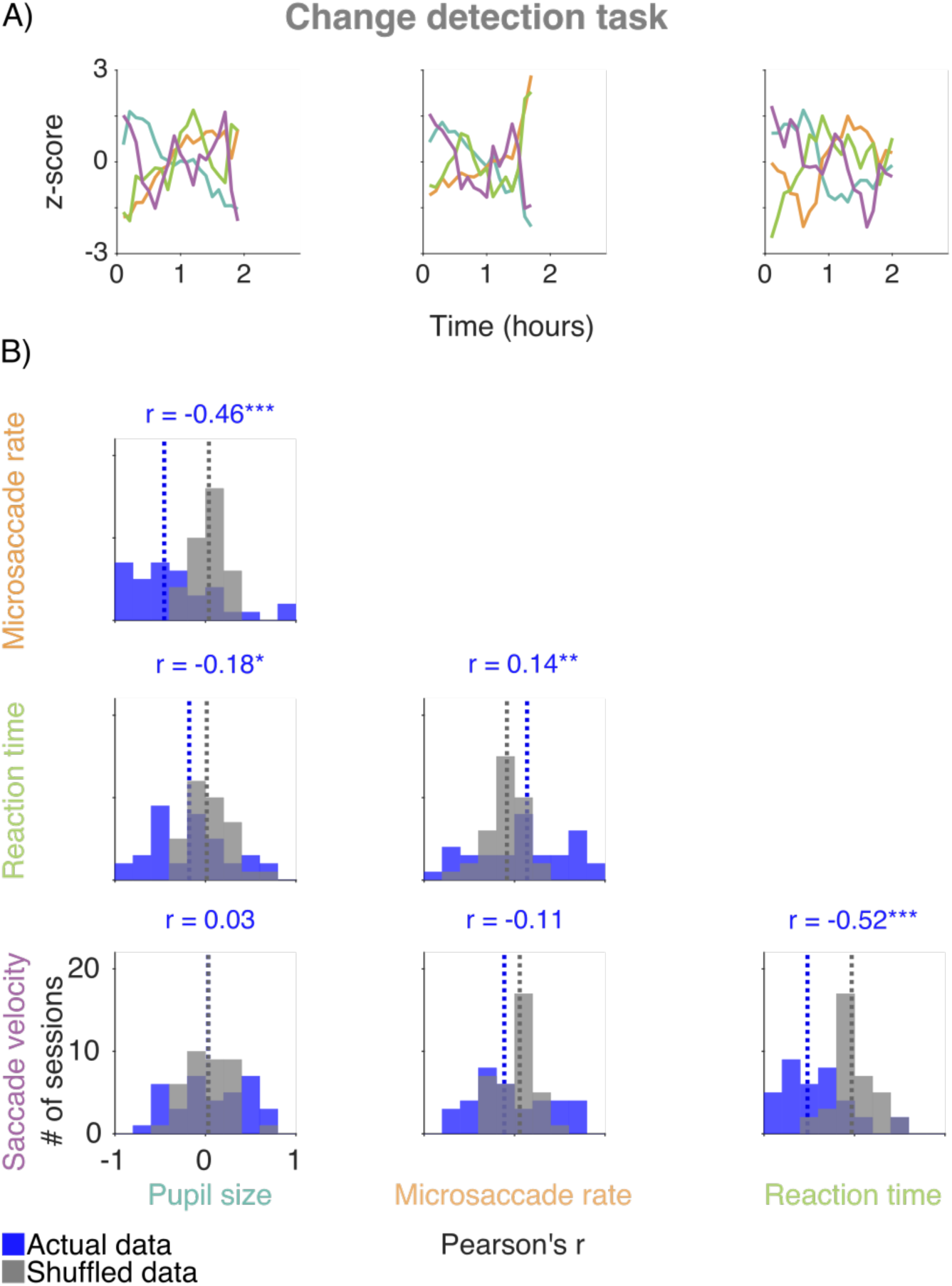
Correlations between the eye metrics on the change detection task. (A) Three example sessions from Monkey 1. (B) Histograms showing actual and shuffled distributions of r values across sessions. Median r values are indicated by dashed lines (blue lines = actual data; gray lines = shuffled data). Actual distributions of r values were compared to shuffled distributions using two-sided permutation tests (difference of medians). p < 0.05*, p < 0.01**, p < 0.001***.

**Figure 3 - figure supplement 2.**
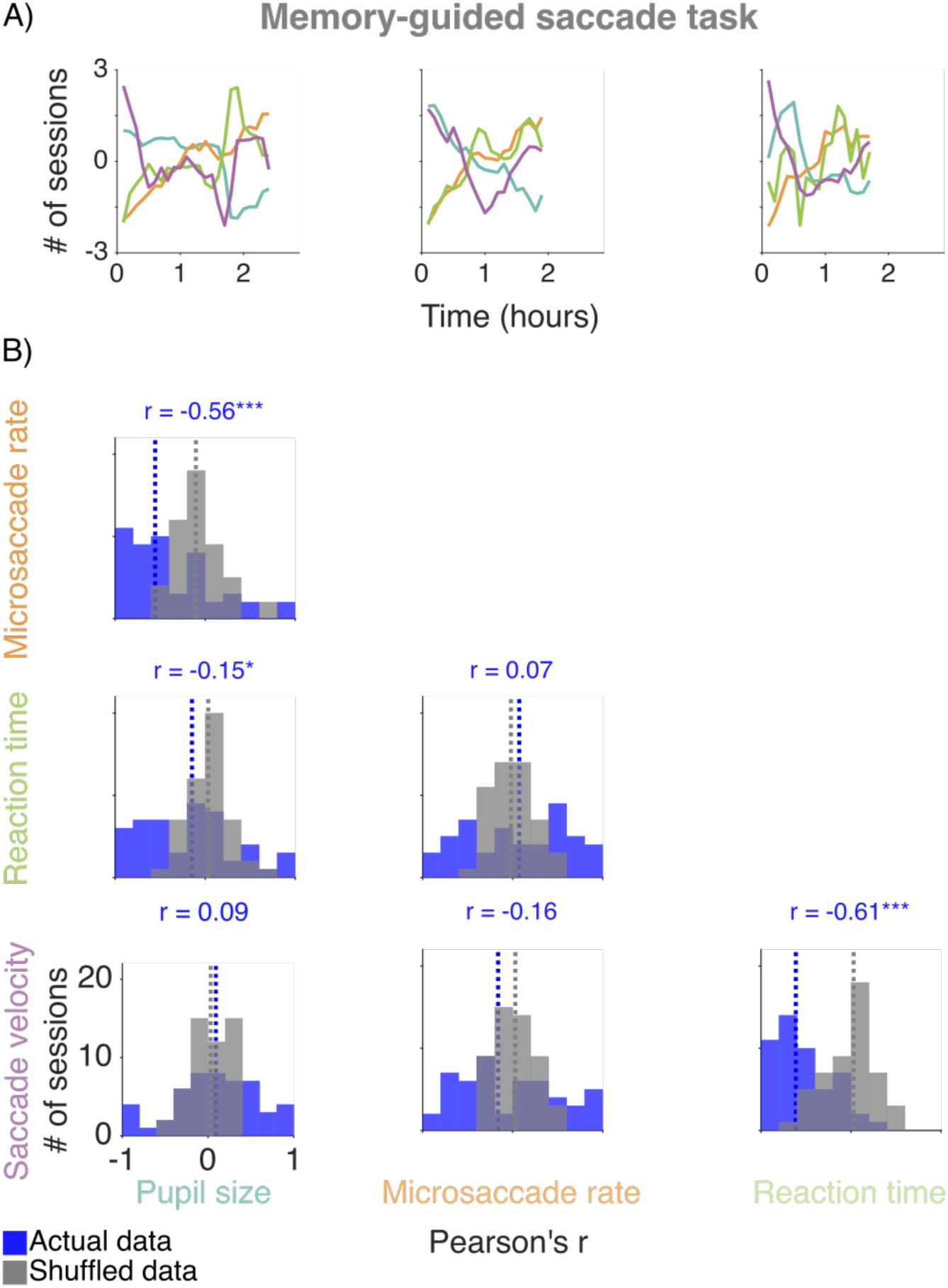
Correlations between the eye metrics on the memory-guided saccade task. (A) Three example sessions from Monkey 1. (B) Histograms showing actual and shuffled distributions of r values. Median r values are indicated by dashed lines (blue lines = actual data; gray lines = shuffled data). Actual distributions of r values were compared to shuffled distributions using two-sided permutation tests (difference of medians). p < 0.05*, p < 0.01**, p < 0.001***.

**Figure 4 - figure supplement 1.**
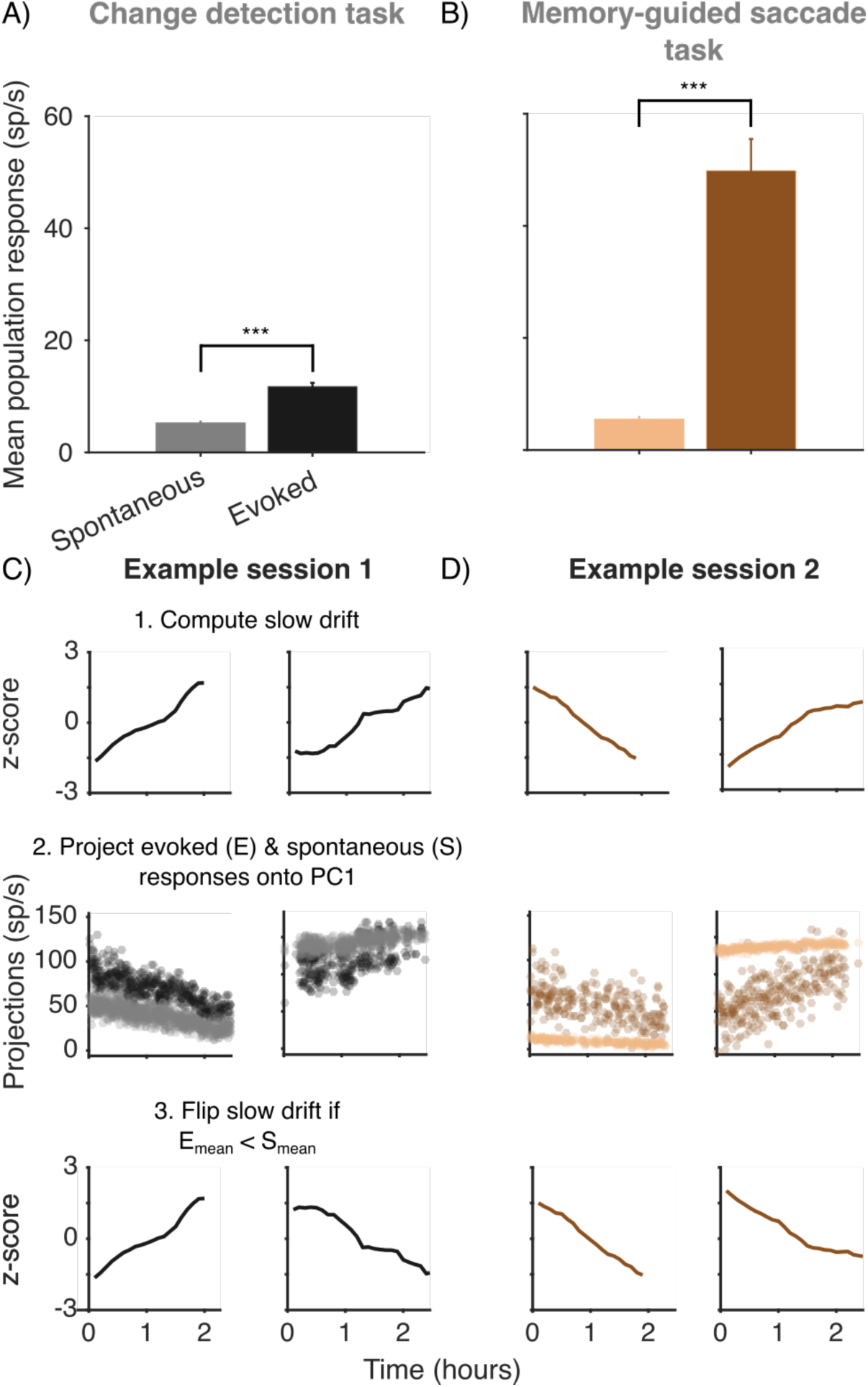
Aligning slow drift across sessions. (A) Bar charts showing the mean spontaneous and evoked population response across sessions on the change detection task. Spontaneous activity was calculated during fixation periods, whereas evoked activity was calculated during stimulus periods. (B) Same as (A) but for the memory-guided saccade task. Spontaneous activity was recorded during the delay period, whereas evoked activity was recorded during the presentation of the target stimulus. In (A) and (B) the mean evoked population response was significantly higher for evoked than spontaneous activity (two-sided permutation tests, difference of means). (C) For each session, slow drift on the change detection task was aligned by projecting evoked and spontaneous responses onto the first principal component. The sign of the slow drift was flipped if the mean projection value for the evoked responses was less than that for the spontaneous responses i.e., if the relationship shown in (A) for unprojected data did not hold true. (D) Same as (C) but for the memory-guided saccade task. p < 0.05*, p < 0.01**, p < 0.001***. Error bars represent ±1 SEM.

**Figure 4 - figure supplement 2.**
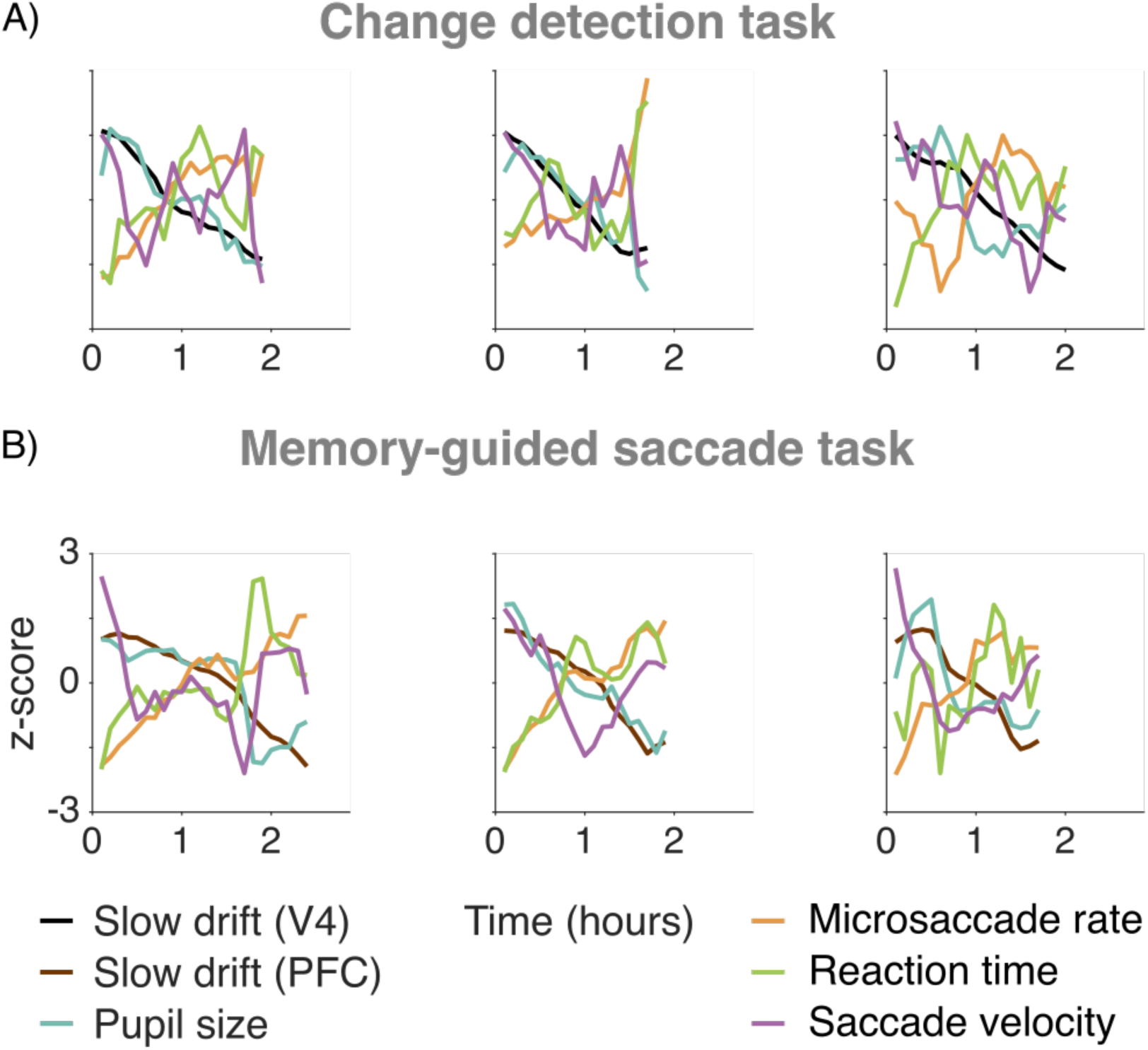
Correlations between slow drift and the eye metrics. (A) Three example sessions from Monkey 1 on the change detection task. (B) Same as (A) but for the memory-guided saccade task (same subject).

**Figure 4 - figure supplement 3.**
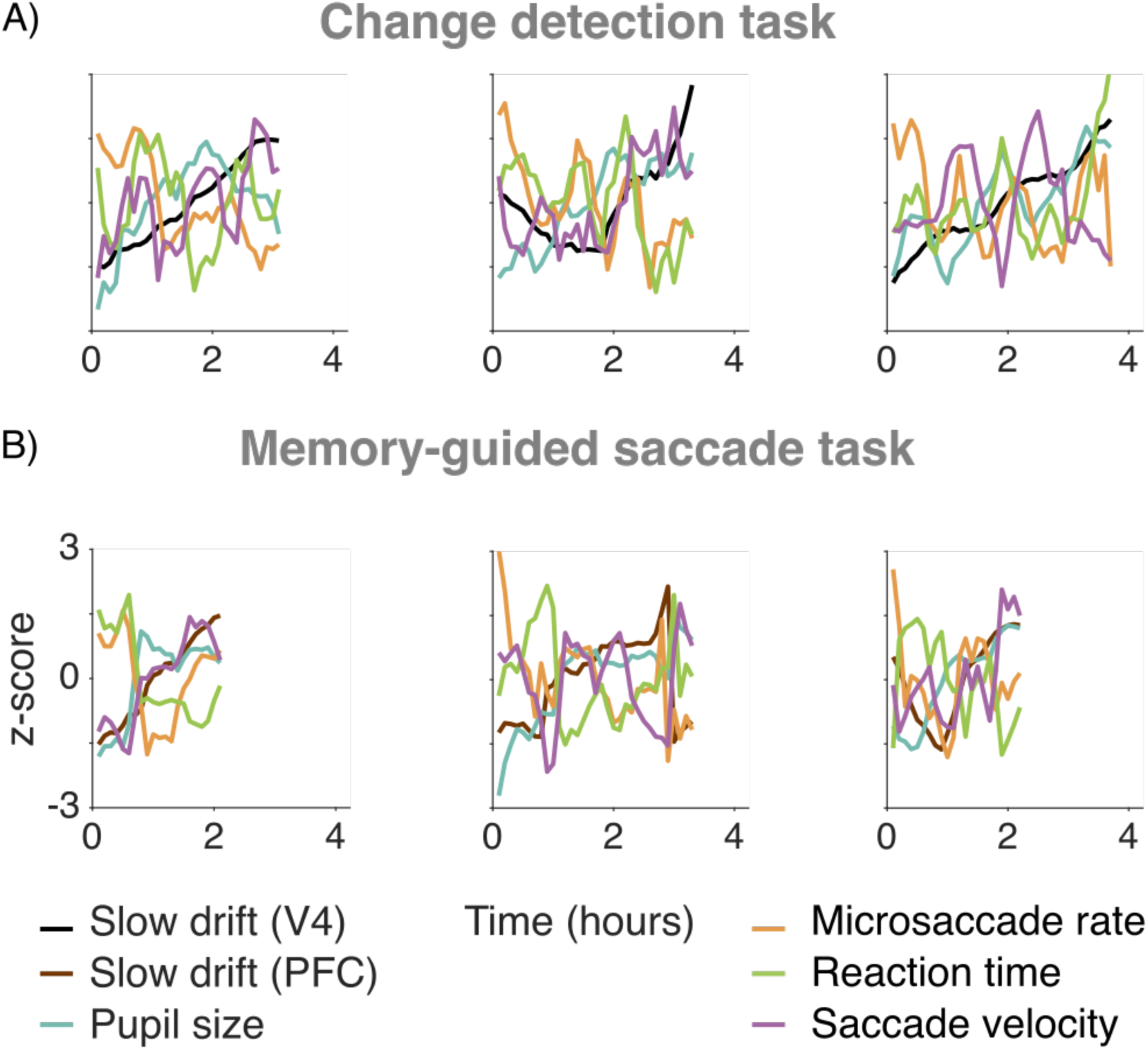
Correlations between slow drift and the eye metrics. (A) Three example sessions from Monkey 2 on the change detection task. (B) Same as (A) but for the memory-guided saccade task (same subject).

**Figure 4 - figure supplement 4.**
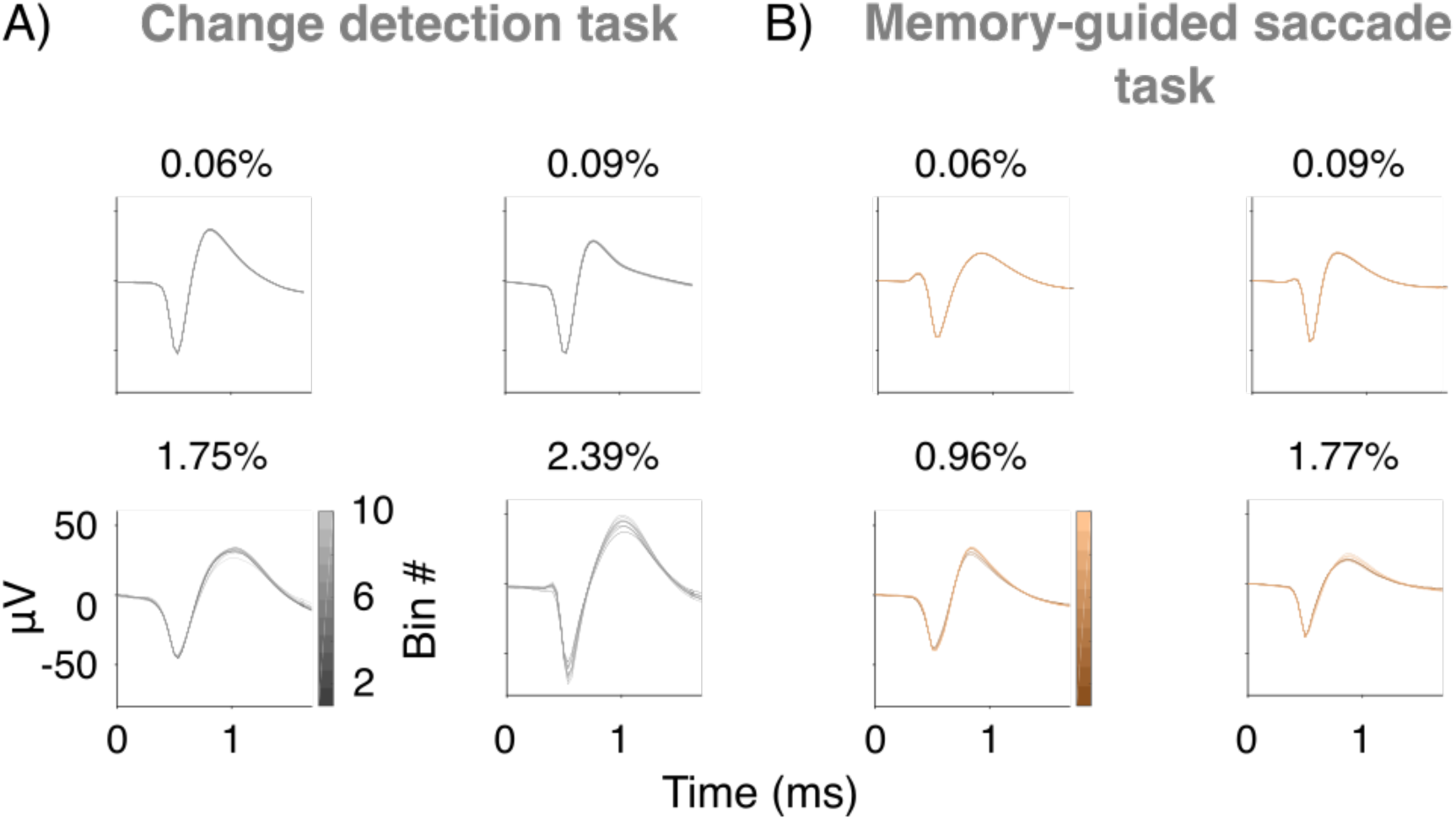
Controlling for neural recording instabilities. (A) Percent waveform variance of four example neurons recorded from Monkey 1 during a single session on the change detection task. (B) Same as (A) but for the memory-guided saccade task (same subject).

**Figure 5 - figure supplement 1.**
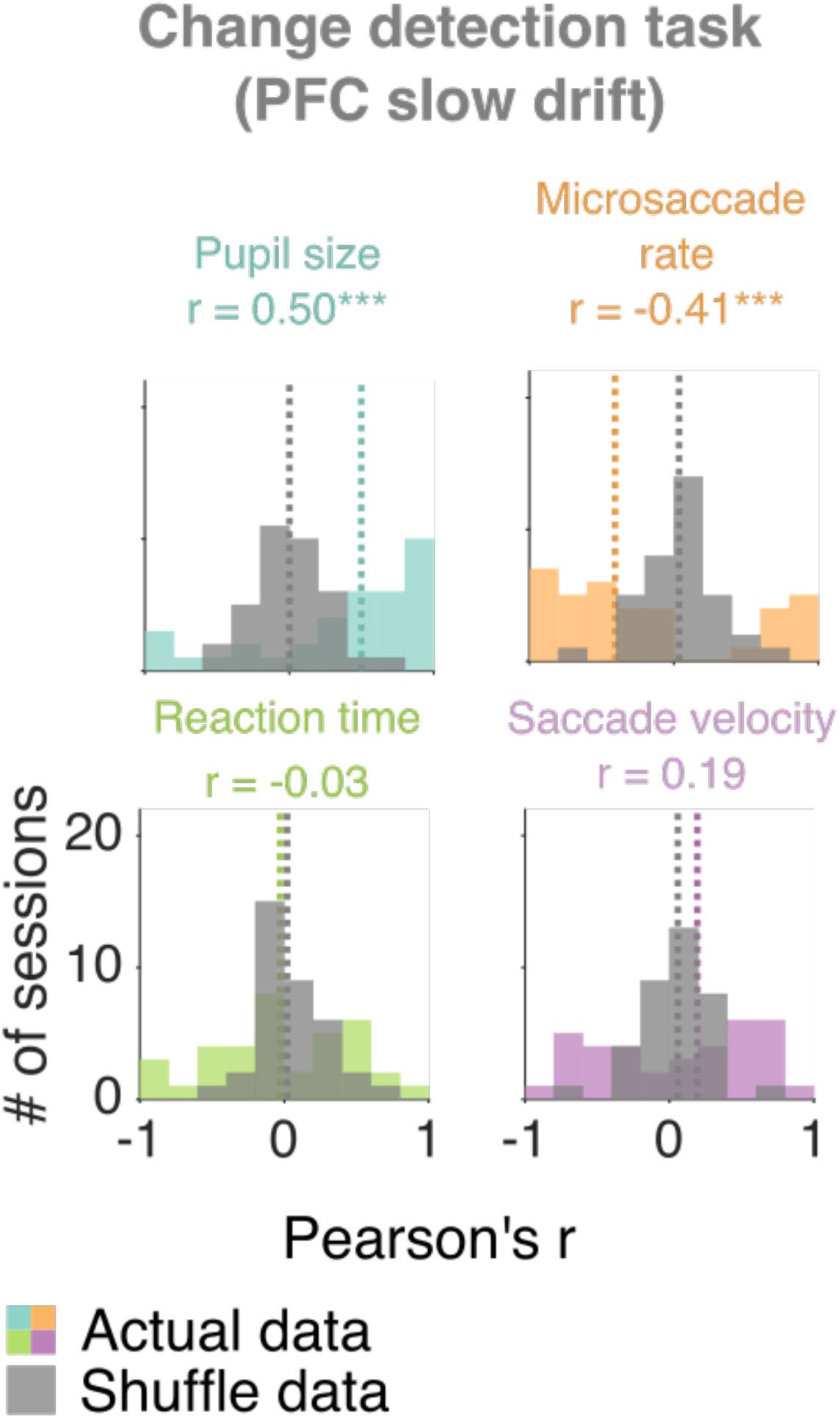
Correlations between the eye metrics and PFC slow drift on the change detection task. Histograms showing actual and shuffled distributions of r values. Median r values across sessions are indicated by dashed lines (colored lines = actual data; gray lines = shuffled data). Actual distributions of r values were compared to shuffled distributions using two-sided permutation tests (difference of medians). p < 0.05*, p < 0.01**, p < 0.001***.

## Acknowledgements

D.I. was supported by National Institutes of Health (NIH) Grant T32 GM-008208 and the ARCS Foundation Thomas-Pittsburgh Chapter Award. M.A.S. was supported by NIH Grants R01 EY-022928, R01 MH-118929, R01 EB-026953, and P30 EY-008098; NSF Grant NCS 1734901; a career development grant and an unrestricted award from Research to Prevent Blindness; and the Eye and Ear Foundation of Pittsburgh. A.C.S. was supported by NIH grant K99EY025768. S.B.K. was supported by NIH Grant T32 EY-017271. The authors would like to thank Ms. Samantha Schmitt for assistance with surgery and data collection.

